# Proteogenomic view of cancer epigenetics: the impact of DNA methylation on the cancer proteome

**DOI:** 10.1101/340760

**Authors:** Majed Mohamed Magzoub, Marcos Prunello, Kevin Brennan, Olivier Gevaert

## Abstract

Aberrant DNA methylation disrupts normal gene expression in cancer and broadly contributes to oncogenesis. We previously developed MethylMix, a model-based algorithmic approach to identify epigenetically regulated driver genes. MethylMix identifies genes where methylation likely executes a functional role by using transcriptomic data to select only methylation events that can be linked to changes in gene expression. However, given that proteins more closely link genotype to phenotype recent high-throughput proteomic data provides an opportunity to more accurately identify functionally relevant abnormal methylation events. Here we present ProteoMix, which refines nominations for epigenetic driver genes by leveraging quantitative high-throughput proteomic data to select only genes where DNA methylation is predictive of protein abundance. Applying our algorithm across three cancer cohorts we find that ProteoMix narrows candidate nominations, where the effect of DNA methylation is often buffered at the protein level. Next, we find that ProteoMix genes are enriched for biological processes involved in cancer including functions involved in epithelial and mesenchymal transition. ProteoMix results are also enriched for tumor markers which are predictive of clinical features like tumor stage and we find clustering on ProteoMix genes captures cancer subtypes.

## Introduction

Genomic characterization can elucidate underlying biology, disease etiology and reveal biomarkers of cancer development and progression; however, each molecular feature is susceptible to different sources of biological and technical measurement noise and provides only one view on the cell state. Therefore, comprehensive studies are needed to understand the molecular basis of disease. Toward this end a multi-institutional consortium, The Cancer Genome Atlas (TCGA), has extensively characterized numerous cancer sites producing genome wide data for mutations, copy number alterations (CNA), RNA expression, microRNA expression, and DNA methylation (1–5). As part of this project, the proteome was initially probed using protein array Reverse Phase Protein Assay (RPPA) technology. However, antibody based analysis are inherently limited because of the reduced coverage and inability to easily compare across proteins due to differential binding effects (6,7). Transcending these limitations, recent advancements in proteomics through high sensitivity mass-spectrometry (MS) are opening new opportunities in cancer research (8). To accelerate the uptake of proteomics the Clinical Proteomic Tumor Analysis Consortium (CPTAC) is performing proteomic analyses of TCGA tumor bio-specimens for a growing number of tissue types and establishing standardized workflows using high-throughput liquid chromatography tandem mass-spectrometry (LC-MS/MS) to capture the proteome as a whole (6,9,10).

To best leverage this new technology comparative analysis between protein abundance and RNA expression can highlight factors influencing concordance and inform how to best interpret proteomic data (11). For example, multiple studies have proven that concordance between mRNA and protein is highly variable, such that one cannot be used to reliably predict the other. Correlation between mRNA and protein has been repeatedly shown to vary by tissue type and cancer status among other molecular features like biological function or molecular stability (7). It was shown across multiple cancers that dynamic proteins involved in metabolism show strong agreement whereas housekeeping proteins and RNA processing proteins are weakly or negatively correlated (6,9,10). So, although many biological functions are regulated primarily through RNA expression – producing moderate correlation between proteomic and transcriptomic data, with mean spearman rho: 0.23 - 0.47 – post-transcriptional mechanisms also play a significant role that cannot be overlooked.

The proteome represents the final link from genotype to molecular phenotype, so proteins are of special importance among molecular features and likely provide a more accurate depiction of cell state; this enhanced view on disease can be leveraged to assess functional effects of upstream aberrations, such as epigenetic modifications. Multi-level epigenetic features such as DNA methylation and histone modification work in concert to regulate gene expression. DNA-methylation, the covalent addition of methyl groups to CpG dinucleotides to form 5-methylcytosine (5mC), is catalyzed by DNA methyltransferases, and is influenced by both environmental and hereditary factors (12). Previous studies have shown that DNA methylation plays a key role in health and is involved in processes of embryonic development and cellular differentiation, where changes can occur through imprinting, inheritance, or de novo events (13,14). Furthermore, DNA methylation has been numerously cited as a potentially causative event in cancer (15,16). Among potential DNA methylation drivers, silencing of tumor suppressors through promoter CpG island hypermethylation is best understood and linked to corresponding gene silencing (13,17,18). Global hypo-methylation on the other hand can potentially result in genomic instability and reactivation of oncogenes (12,13,15).

To elucidate the role of DNA methylation in disease, our goal is to investigate whether linking proteomic data with DNA methylation data identifies key genes, describes molecular features and subtypes in cancer. Previously we presented MethylMix an algorithm that formalizes the identification of DNA methylation driver genes using a model-based approach (19–23). Recognizing the complex role of the methylome in epigenetic regulation of cancer, MethylMix uses mRNA data to select only differentially methylated genes that show down-stream effect on gene expression. This selects for likely functional aberrations with the aim of discriminating between true driver genes, and passenger events which are characteristic of genome wide dysfunction in cancer. Herein we present ProteoMix which refines candidate nominations for epigenetic driver genes by excluding aberrations that are buffered at the protein level; this likely selects for events which are functional over those which may accumulate during cancer but do not drive pathogenesis. Using quantitative MS data from three cancer cohorts: breast invasive carcinoma, colorectal adenocarcinoma, and ovarian serous cystadenocarcinoma, we report ProteoMix’ gene identifications, which include potential markers and therapeutic targets. We describe ProteoMix’ ability to elucidate key molecular and higher level disease features and evaluate ProteoMix’ performance against MethylMix. In summary, our study highlights the differences between integrated epigenomic-proteomics and epigenomic-transcriptomics analyses.

## Results

We applied ProteoMix and MethylMix (19–22) across three cancer types with both transcriptomic and proteomic data (Table 1): breast invasive carcinoma (BRCA), colorectal adenocarcinoma (COADREAD), and ovarian serous cystadenocarcinoma (OV). Our analysis compares genes identified by ProteoMix and MethylMix (Supplementary Table 1), specifically examining the biological and clinical relevance of each model’s output and utility for downstream analysis.

**Table 1.** Overview of number of genes, CpG Clusters, and samples used for each TCGA cancer site analysis.

|  | N Genes | N CpG Clusters | N Samples: Gene<br>Expression &<br>Protein<br>Abundance | N Samples: Tumor<br>Tissue Methylation | N Samples: Normal<br>Tissue Methylation |
| --- | --- | --- | --- | --- | --- |
| BRCA | 2514 | 3693 | 78 | 972 | 123 |
| COADREAD | 2848 | 4288 | 85 | 614 | 78 |
| OV | 1896 | 3693 | 168 | 582 | 8 |

### ProteoMix narrows candidate nominations for epigenetically driven genes

For each cohort both models identify genes that are 1) differentially methylated when compared to normal adjacent tissue and 2) functionally predictive of downstream effects at the level of RNA expression in the case of MethylMix or protein abundance in the case of ProteoMix (Figure 1). Among all three cancer cohorts we observe significant correlations between RNA expression and protein abundance (mean rho: 0.23-0.47), indicating that most genes are regulated at the transcript level (Supplementary Table 2). Therefore, it is unsurprising that ProteoMix shows high agreement with MethylMix, where more than 90% of ProteoMix genes are also identified by MethylMix. However, ProteoMix lists are more conservative identifying fewer candidate genes across all three cancers, where often the effect of methylation is present at the RNA level, but not detected at the protein level (Figure 1), likely because they are buffered at the protein due to post-transcriptional, translational, or degradation regulation. Therefore, ProteoMix better enriches for methylation-states that more likely execute functional roles in cancer development.

**Figure 1:**
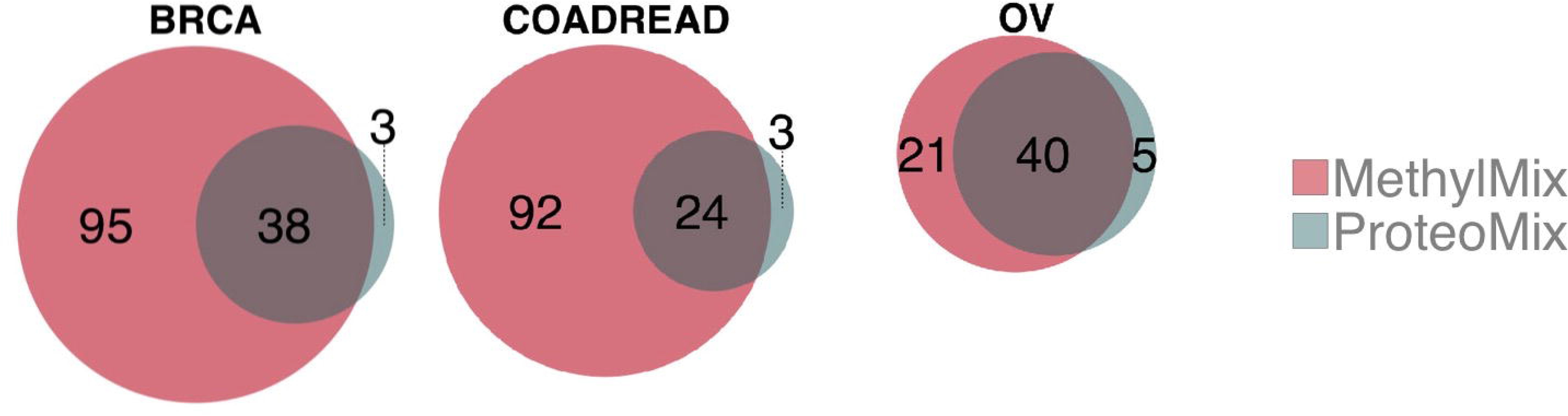
Venn diagrams comparing the number of reported genes that are differentially methylated and functionally predictive for MethylMix and ProteoMix.

### ProteoMix identifies new genes with significant methylation effects only at the protein level

For each cancer cohort ProteoMix also identifies a few unique driver genes, the majority of which have documented roles in carcinogenesis. Explanative mechanisms by which the effect of DNA methylation may be undetected at the RNA level but functional at the protein level are further addressed below in the discussion.

In breast cancer ProteoMix discovers three novel differentially methylated genes of diverse biological functions. ProteoMix detects a functional effect of hypo-methylation in the untranslated region (UTR) of *EHF*, which is a well-studied transcription factor involved in HER2 mediated epithelial differentiation (35); a likely oncogene, knockdown of *EHF* has been shown to inhibit tumor invasion and proliferation (36). Next, ProteoMix identifies hyper-methylation of *FSTL1*, an autoantigen that promotes immune response. This candidate tumor suppressor, *FSTL1*, has also been shown to mediate tumor immune evasion in nasopharyngeal cancer through hyper-methylation silencing (37). ProteoMix also reports hyper-methylation of *DHX40* which has an unclear link to cancer; although it is of note that RNA splicing proteins – like *DHX40* – are highly stable, perhaps explaining the particularly stronger effect of DNA methylation on protein abundance than mRNA (38) (Supplementary Table 1).

In colorectal cancer ProteoMix recovers several genes associated with immune function and inflammation, which is known to play a key role in pathogenesis. ProteoMix uniquely identifies a functional effect of UTR hypo-methylation of the *PTPRC* gene. *PTPRC* belongs to a family of protein tyrosine phosphatase which contains oncogenes regulating cell growth and differentiation. *PTPRC* is also related to tumor necrosis and disrupts normal T- and B-cell signaling through SRC kinase pathways - which are separately implicated in colorectal cancer through amplification (9,39). Next, ProteoMix identifies upregulation of *S100A9* through promoter hypo-methylation. Of note, elevated *S100A9* mRNA and protein levels are commonly observed in many conditions associated with inflammation (40); additionally in hydropharangeal cancer where knockdown inhibited cell growth and invasion, *S100A9* is also prognostic of worse outcome and indications like metastasis (41). Of note ProteoMix filtered out functional effects of a UTR hypo-methylation in *S100A9* previously detected by MethylMix. Next, ProteoMix identifies hyper-methylation across the promoter region of *LTF*, a likely tumor suppressor which is produced by neutrophils to regulate growth and differentiation. In the context of colorectal tissue *LTF* has been shown to restrict inflammation by regulating T cell interaction (42). Additionally, gene expression of *LTF* has previously been shown to correlate with tumor size and survival in breast cancer (43). Lastly, ProteoMix uniquely identifies hypo-methylation mediated upregulation of *DAK*, also known as *TKFC*, which is related to virus-associated chronic inflammation (44).

ProteoMix picks up hypo-methylation states in five new genes in ovarian cancer related to processes of invasion and proliferation. ProteoMix uniquely identifies hypo-methylation in the promoter region of *EVL* a key regulator of the actin cytoskeleton, associated with invasion and metastasis. Overexpression of *EVL* is also indicative of advanced stage in breast cancer (45) and has been implicated in malignancies due to inappropriate recombination (46). ProteoMix discovers elevated TSTA3 expression caused by gene body hypo-methylation. *TSTA3* is linked to malignant transformations through abnormal glycosylation and controls cell proliferation and invasion by regulating CXCR4 chemokine mediated T cell signaling; additionally in breast cancer, high *TSTA3* expression correlated with poor survival (47). Next, ProteoMix detects promoter hypo-methylation mediated upregulation of *HMGB3*, a well-documented oncogene implicated in several cancers including breast, lung, esophageal, bladder, and colorectal cancers (48–51). *HMGB3* promotes cell proliferation in colorectal cancer cells through regulation of *MYC*, where genes from same family have been linked to decreased *MYC* expression which is unique to proliferative subtypes of ovarian cancer (26). Lastly, ProteoMix also identifies hypo-methylation in two mitochondrial genes *ATP5D* and *SPG7*, speculatively linked to cancer through metabolic function (52).

### ProteoMix genes are enriched for biological processes involved in cancer

We conducted enrichment analysis to identify biological processes that are overrepresented in ProteoMix and MethylMix genes (Supplementary Table 3). Given the large proportion of common genes, across all three cancers both models capture many of the same annotations. However, comparing enrichments found for each cancer site, we find that broadly ProteoMix results include more significant enrichments for functions associated with cell adhesion and migration of epithelial and endothelial cells; these processes increase cell motility and invasiveness and are indicative of epithelial to mesenchymal transition (EMT) which is key to cancer development. Additionally, we observed that enrichment for immune functions are highly variable between each model’s results.

Comparing unique annotations among breast cancer genes, ProteoMix includes enrichments for responses to growth factor, angiogenic processes, and immune development, whereas MethylMix uniquely captures specific immune processes associated with inflammation and migration of T cells and leukocytes. However, the MethylMix gene list is also enriched for homeostasis, metabolic, and several other functions with no clear relevance to cancer including taste perception and skin and limb morphogenesis. Strikingly in colorectal cancer, although the ProteoMix gene list is shorter it captures all MethylMix enrichments and adds numerous new annotations including EMT related functions like cell migration, cell adhesion, and mesenchyme development. ProteoMix also uniquely enriches for: 1) immune response, toll-like receptor recognition, and cellular functions of immunoglobulin, B cells, leukocytes, lymphocytes and mucosal-associated lymphoid tissue; 2) signaling functions mediated through NF-kappaB and integrin; 3) and other functions include intra-cellular transport and hormone secretion. For ovarian cancer, the ProteoMix genes are uniquely enriched for negative regulation of B cell differentiation, but misses many immune response functions involving cytokine production and mast cells only captured by MethylMix, which also uniquely captures endothelial cell proliferation.

Looking for commonalities between cancers we find more shared enriched biological processes when comparing among ProteoMix results, suggesting that ProteoMix better captures the underlying similarities in disease etiology. Comparing breast and ovarian cancer we find both lists share enrichments for processes involved in apoptotic signaling. Each gene list however captures different aspects of immune response, with ProteoMix identifying common enrichments of B cell differentiation and MethylMix uniquely capturing leukocyte migration and inflammation response. MethylMix also uniquely identifies processes regulating cellular motility and migration, but also identifies commonalities in hormone metabolism and other metabolic processes. When comparing across breast and colorectal cancers MethylMix identifies no shared biological processes, whereas ProteoMix finds common enrichments for cell migration, exocytosis, and hormone levels. Lastly looking at ovarian and colorectal cancers, both sets of ProteoMix genes are enriched for several immune response related processes including leukocyte, myeloid, and granulocyte differentiation. Both lists also share enrichments for regulation of cell death and other processes including protein transport and xenobiotic metabolism, which is possibly associated with platinum sensitivity (53).

### ProteoMix genes are enriched for tumor progression markers

Taking an orthogonal approach, we identified putative biomarker of disease progression based on correlations between gene expression and clinical features (Table 2). Although ProteoMix gene lists contain much fewer identifications we find that across all three cancers that ProteoMix’ lists include a larger proportion of markers of tumor stage and size and show stronger odds of containing such genes (Table 2). The greatest difference in frequency of tumor stage marker is observed in breast cancer where 12% versus 7% of genes show correlation in ProteoMix and MethylMix gene lists respectively. The most significant associations however are observed in colorectal cancer where 15% of ProteoMix genes show correlation between gene expression and tumor stage, this includes LTF which is mentioned among unique ProtoMix genes (Table 2A, Supplementary Table 1). The same trend applies when correlating gene expression with tumor size where the largest difference in enrichment can be seen in colorectal cancer where 7% versus 3% of genes correlate with size when comparing models. However, the enrichment is much stronger for breast cancer where 29% of genes correlate with tumor size compared to 21% of MethylMix genes (Table 2B).

**Table 2.**
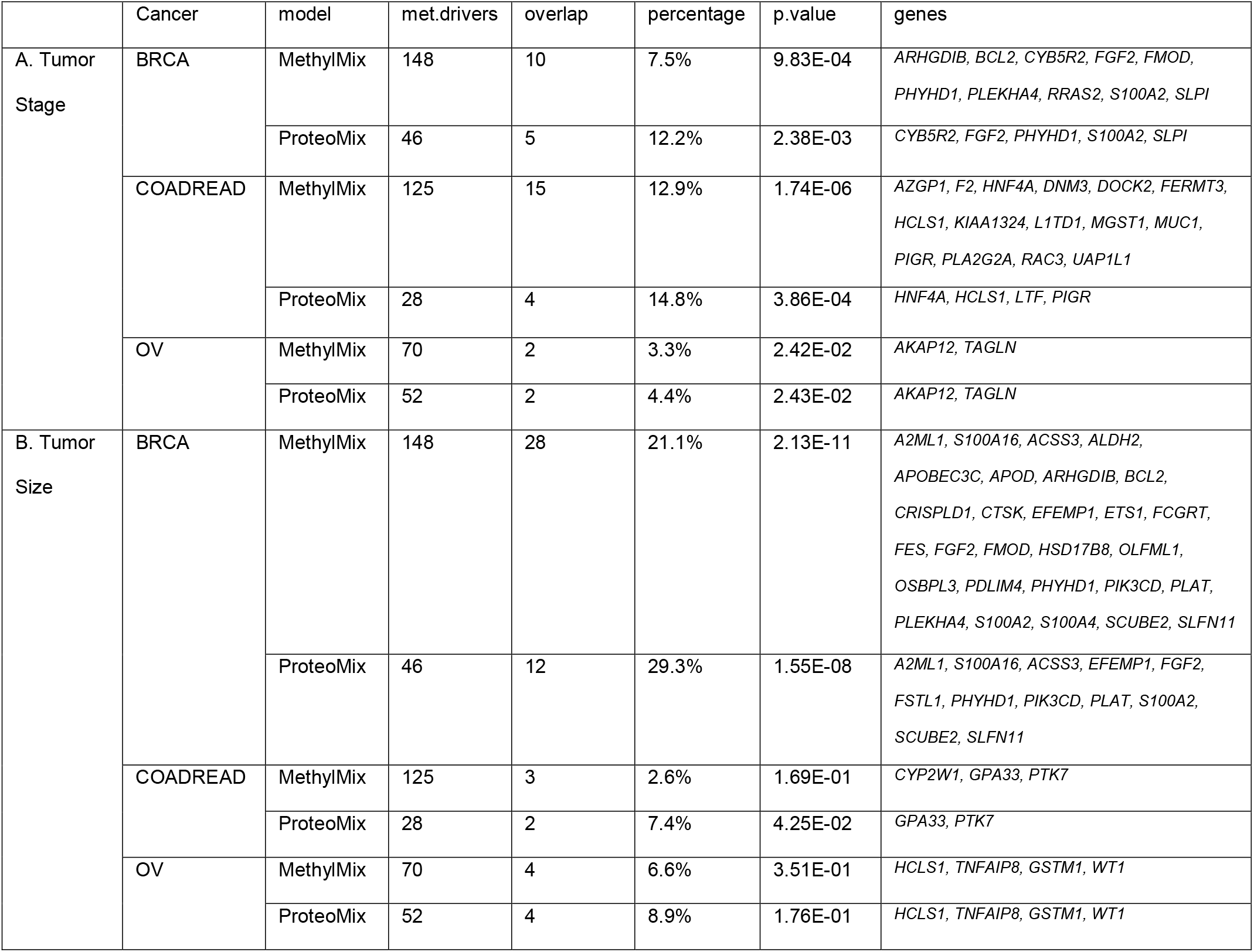
Report of overlap between MethylMix and ProteoMix genes with tumor progression markers produced using a fisher exact test.

|  | Cancer | model | met.drivers | overlap | percentage | p.value | genes |
| --- | --- | --- | --- | --- | --- | --- | --- |
| A. Tumor Stage | BRCA | MethylMix | 148 | 10 | 7.5% | 9.83E-04 | *ARHGDIB, BCL2, CYB5R2, FGF2, FMOD, PHYHD1, PLEKHA4, RRAS2, S100A2, SLPI* |
|  |  | ProteoMix | 46 | 5 | 12.2% | 2.38E-03 | *CYB5R2, FGF2, PHYHD1, S100A2, SLPI* |
|  | COADREAD | MethylMix | 125 | 15 | 12.9% | 1.74E-06 | *AZGP1, F2, HNF4A, DNM3, DOCK2, FERMT3, HCLS1, KIAA1324, L1TD1, MGST1, MUC1, PIGR, PLA2G2A, RAC3, UAP1L1* |
|  |  | ProteoMix | 28 | 4 | 14.8% | 3.86E-04 | *HNF4A, HCLS1, LTF, PIGR* |
|  | OV | MethylMix | 70 | 2 | 3.3% | 2.42E-02 | *AKAP12, TAGLN* |
|  |  | ProteoMix | 52 | 2 | 4.4% | 2.43E-02 | *AKAP12, TAGLN* |
| B. Tumor Size | BRCA | MethylMix | 148 | 28 | 21.1% | 2.13E-11 | *A2ML1, S100A16, ACSS3, ALDH2, APOBEC3C, APOD, ARHGDIB, BCL2, CRISPLD1, CTSK, EFEMP1, ETS1, FCGRT, FES, FGF2, FMOD, HSD17B8, OLFML1, OSBPL3, PDLIM4, PHYHD1, PIK3CD, PLAT, PLEKHA4, S100A2, S100A4, SCUBE2, SLFN11* |
|  |  | ProteoMix | 46 | 12 | 29.3% | 1.55E-08 | *A2ML1, S100A16, ACSS3, EFEMP1, FGF2, FSTL1, PHYHD1, PIK3CD, PLAT, S100A2, SCUBE2, SLFN11* |
|  | COADREAD | MethylMix | 125 | 3 | 2.6% | 1.69E-01 | *CYP2W1, GPA33, PTK7* |
|  |  | ProteoMix | 28 | 2 | 7.4% | 4.25E-02 | *GPA33, PTK7* |
|  | OV | MethylMix | 70 | 4 | 6.6% | 3.51E-01 | *HCLS1, TNFAIP8, GSTM1, WT1* |
|  |  | ProteoMix | 52 | 4 | 8.9% | 1.76E-01 | *HCLS1, TNFAIP8, GSTM1, WT1* |

### Clustering on ProteoMix genes captures cancer subtypes

Clustering on methylation has been shown to stratify patients into clinically relevant subgroups (2,20,21,23). We performed consensus clustering using the DM values for ProteoMix and MethylMix genes evaluating clusters sizes from two to six (Table 3); for clarity we discuss clusters at K=2, examining the gross differences between MethylMix and ProteoMix. We evaluated if these epigenetically defined subgroups correspond to previously published subtypes and clinical and genetic features and found that ProteoMix identifies subgroups of patients that enriched for specific cancer subtypes and other molecular features and performs similarly to MethylMix (Supplementary Table 4).

**Table 3.** Summary statistics from consensus clustering analysis across K=2-6 for each cancer; we report inter- and intra-cluster scores along with PAC score.

|  |  | MethylMix |  |  | ProteoMix |  |  |
| --- | --- | --- | --- | --- | --- | --- | --- |
| Cancer | K | Intra | Inter | PAC | Intra | Inter | PAC |
| BRCA | 2 | 97.9 | 29.2 | 0.040 | 99.0 | 27.4 | 0.026 |
|  | 3 | 94.6 | 20.0 | 0.199 | 78.4 | 24.2 | 0.409 |
|  | 4 | 87.0 | 15.7 | 0.206 | 80.0 | 17.6 | 0.381 |
|  | 5 | 76.8 | 13.8 | 0.284 | 68.6 | 14.6 | 0.333 |
|  | 6 | 76.2 | 11.9 | 0.275 | 66.5 | 12.9 | 0.309 |
| COADREAD | 2 | 97.9 | 26.6 | 0.074 | 92.3 | 29.5 | 0.273 |
|  | 3 | 68.9 | 26.0 | 0.570 | 78.4 | 23.8 | 0.502 |
|  | 4 | 64.4 | 20.2 | 0.491 | 69.9 | 19.9 | 0.459 |
|  | 5 | 68.9 | 15.2 | 0.401 | 71.6 | 15.5 | 0.405 |
|  | 6 | 71.6 | 12.1 | 0.305 | 67.8 | 13.2 | 0.343 |
| OV | 2 | 94.0 | 29.0 | 0.180 | 93.9 | 29.2 | 0.134 |
|  | 3 | 81.0 | 23.8 | 0.477 | 80.7 | 23.7 | 0.488 |
|  | 4 | 69.1 | 19.3 | 0.438 | 70.8 | 19.5 | 0.445 |
|  | 5 | 74.1 | 14.6 | 0.349 | 73.6 | 14.7 | 0.337 |
|  | 6 | 67.9 | 12.8 | 0.311 | 73.7 | 12.1 | 0.291 |

In breast cancer ProteoMix clusters significantly correlate with molecular subtypes and other molecular features such as Progesterone and Estrogen Receptor (PR, ER) status (Figure 2A). Similar to other studies our clusters differentiate between canonical breast cancer molecular subtypes: Cluster-1 contains about 90% of patients with Luminal A/B type tumors. Cluster-2 contains 97% of patients with Basal-like tumors and as expected it is enriched for samples negative for ER, PR, or HER2. HER2 and Normal subtypes are less clearly distinguished in ProteoMix clusters but can be found in greater frequency in cluster-2. Among colorectal samples we are able to confirm the CpG island methylator phenotype (CIMP) (Figure 2B). Cluster-1 contains 97% of patients labeled CIMP-High using methylation signatures and 82% of patients labeled Microsatellite Instable/CIMP using gene-expression signatures. The CIMP subtype has known association with MLH1 silencing through hyper-methylation, which is reflected in our ProteoMix subtypes where we find cluster-2 to include the majority of samples with non-silenced MLH1. ProteoMix subtypes also significantly correlate with Microsatellite Instability where samples labeled as Microsatellite Instability-Low (MSI-L) or Microsatellite Stable (MSS) are found by majority in cluster-2. Examining subtypes in ovarian cancer our ProteoMix clusters agree well with molecular subtypes and are significantly correlated (Figure 2C). About half of cluster-1 is comprised of patients labeled as Proliferative, while cluster-2 contains 75% of Immunoreactive subtype and 80% of Differentiated subtype patients, lastly Mesenchymal subtype patients can be found with relatively equal frequencies in each cluster (54–56). ProteoMix clusters also significantly correlate with tumor features, where cluster-1 and cluster-2 roughly correspond patients with lower-grade and higher-grade tumors.

**Figure 2:**
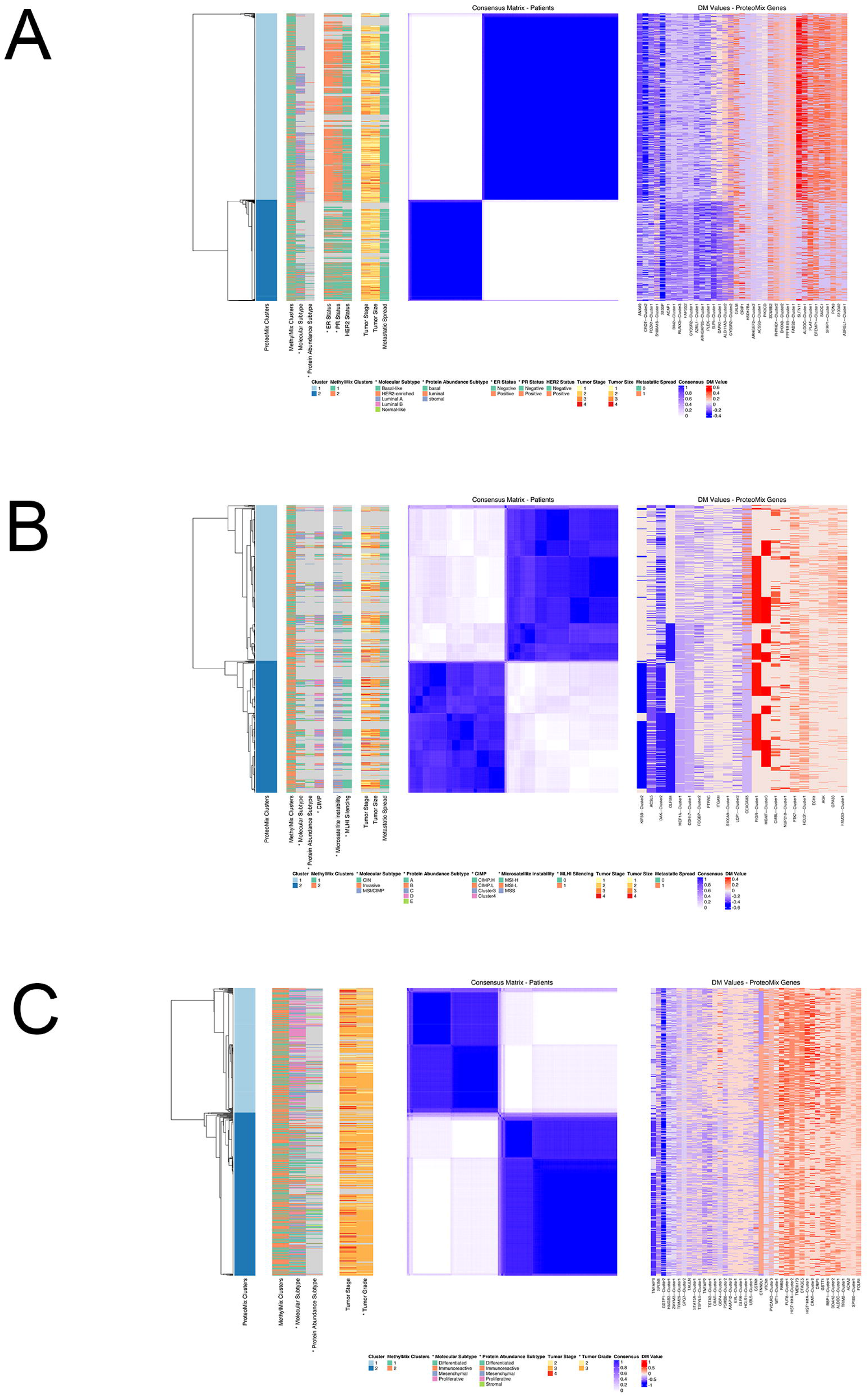
Consensus clustering and methylation profiles for three cancer sites at K=2. (A) breast cancer (BRCA); colorectal cancer (COADREAD); ovarian cancer (OV). Middle panels: visualization of the consensus clustering with blue indicating high consensus and white indicating low consensus. Right panels: methylation profile with red indicating hyper-methylation, white indicating normal methylation, and blue indicating hypo-methylation. Left panels: visualization of additional molecular and clinical features. Non-reported values are marked in grey. Statistically significant overlaps, found using Chi-squared and Kruskal-Wallis tests, are marked with asterisks.

## Discussion

Epigenetic aberrations contribute to oncogenesis, where DNA hypermethylation inactivates tumor suppressor genes, while hypomethylation is known to promote genomic instability and activate oncogenes (12,20). Therefore, DNA methylation has potential to inform patient treatment and improve patient outcomes through new diagnostics and therapeutics. When identifying epigenetically driven cancer genes, it is of note that most biological functions – subject to genomic and epigenomic dysregulation – are ultimately executed at the protein level, so we can expect neutralization of non-functional upstream effects at - or before - the proteome. Herein we confirm the potential of using proteomic data to elucidate functional DNA methylation events by conducting the first genome wide analysis of epigenome-proteome relationships across three large human cancer cohorts. We present ProteoMix, a data-driven model which formalizes the identification of abnormally methylated genes that are predictive of protein abundance ProteoMix, like MethylMix, uses a model-based approach, negating the use of arbitrary user-defined thresholds for abnormal DNA methylation, and identifies subpopulations of hypo or hypermethylated samples within a heterogeneous population. By integrating DNA methylation array and quantitative MS technologies, ProteoMix identifies candidate epigenetic driver genes with clinical value as potential therapeutic targets and protein biomarkers for assessing prognosis and treatment stratification. ProteoMix builds on our model MethylMix and addresses the potential limited predictive value of mRNA as proxy for phenotype due to the role of post-transcriptional mechanisms.

ProteoMix identifies oncogenes and tumor suppressors and – with the exception of a few genes – returns a subset of MethylMix identifications, where often the effect of DNA Methylation does not propagate to the proteome (Figure 1, Supplementary Table 1). In other cancer studies similar buffering has been observed in both cis and trans CNA effects, suggesting that many detectable aberrations in cancer do not manifest in expression changes at the protein level (6,10). Otherwise put, many abnormally methylated genes are likely only passengers and do not functionally contribute to cancer development. Identification of a reduced set of genes in our study has pragmatic benefits for cancer research, where narrowing nominations to fewer high-quality candidates increases the likelihood of finding true targets; strongest candidates include genes identified by both models that show negative correlation between DNA methylation and both gene expression and protein abundance, and therefore have clear biological interpretations amenable to validation in the laboratory. Similar methods to identify true targets have been described, where genes that show correlation between mRNA and protein are more likely to have tumor promoting effects (10). Conversely, novel ProteoMix identifications should be taken with due consideration given the lack of clear mechanisms explaining how changes in DNA methylation may alter protein levels, but be undetectable at the transcript level - plausible explanations that remain to be tested include erroneous or noisy gene expression data, low mRNA stability or alternative splicing confounding expression at the RNA level. Nevertheless, most new identifications are well supported to have tumor promoting effects and therefore warrant further investigation to uncover how DNA-methylation may influence regulation of genes like *EHF, FSTL1, PTPRC, S100A9, LTF, EVL*, and *TSTA3*. Importantly, in all these cases the type of DNA methylation is consistent with gene function, where known tumor-suppressors are hyper-methylated and oncogenes are hypo-methylated at regions where DNA methylation negatively regulates transcription.

Taken together ProteoMix genes highlight important features in cancer related to tumor features and subtypes, additionally ProteoMix captures oncogenic biological processes. Using enrichment analysis ProteoMix identifies key aspects of cancer development such as processes related to angiogenesis, EMT, immune function, and proliferative signaling (Supplementary Table 3). ProteoMix also elucidates more shared annotations between cancer types, and thus a greater ability to identify genes of core cancer pathways that are shared across cancer sites. Next, using a completely orthogonal approach we also find that ProteoMix is more descriptive of tumor progression; although our new model produces a reduced number of identifications, ProteoMix genes are more likely to correlate in expression with disease features such as tumor stage and size (Table 2). Lastly, we find ProteoMix performs reasonably well for patient clustering recapitulating established molecular subtypes. Given the limitations of our study, we expect our clustering to have reduced discriminative power, since we limit our observations to genes for which we have both matched gene expression and protein abundance measurements in our analysis and significantly diminish the feature space we used for learning. Nevertheless, we find that ProteoMix performs similarly to MethylMix in identifying cancer subtypes such as Luminal and Basal types of breast cancer, the CIMP type in colorectal cancer and all subtypes in ovarian cancer, with the exception of the Mesenchymal subtype which is the least clearly defined subtype (54) (Figure 2, Supplementary Table 4). These findings suggest the reduced number ProteoMix genes capture the major sources of variation in each cancer cohort and facilitate translatability into feasible panels for testing.

Overall ProteoMix shows practical utility for improving nominations of cancer driver genes and elucidating new mechanisms of cancer development missed by our previous model. More broadly our study supports using proteomic data to better understand how epigenetic deregulation promotes cancer. Similar approaches have been applied and found to potentially improve aspects of patient care. For example, a retrospective analysis of outcomes in an oncology trial for glioblastoma – which tested efficacy of different temozolomide regiments – found that updating the clustering model to incorporate MGMT protein expression and c-MET protein abundance provided better separation of overall survival prognostic groups than incorporating MGMT promoter methylation alone (57). These findings and ours support the claim that protein data combined with DNA methylation is a better way to stratify patients and understand cancer features then using DNA methylation alone.

Although milestone initiatives like TCGA and CPTAC provide valuable date for the acceleration of discovery and research in cancer, we acknowledge the limitations of this study and further work required. A barrier to translation, the number of specimens used here is insufficient to draw conclusive clinical correlations and require replication of these results by independent studies. Importantly molecular measurements used here are also subject to sources of technical and biological bias. For example, it is known that bulk measurements obscure the complex nature of tumor microenvironment which includes many cell types including vascular, lymphatic, and immune cells. This confounding effect is compounded considering that each molecular feature was measured using different tumor fragments, which may have very different cellular compositions due to intra-tumor heterogeneity. Additionally, we recognize further characterization of genome wide proteomic studies is required to fully understand possible biases, such as worse detection of highly hydrophobic and hydrophilic peptides, or low-abundance peptides co-eluting with very high-abundance peptide (9). Moreover, early proteomic techniques such as those utilized in CPTAC’s Common Data Analysis Pipeline have not yet reached the genome level resolution of other omic measurements; these methods require refinement to address low coverage due to inherent limitations of proteolytic measurements such immeasurable peptides that are excessively large or small tryptic fragments and the inability to distinguish some amino acids (9). This reduced coverage to a few thousand genes in our study excludes many genes with possible roles in cancer.

The complex nature of disease development and interplay between interacting biological aberrations – genetic, epigenetic, somatic or germline - often makes it difficult to elucidate causal mechanisms of cancer development’. Furthermore, there is still much work in multi-omics to elucidate causal flows of information influencing cellular physiology and pathology and to discriminate how separate phenomena are linked to create cancer (3,5,56,58). However, integrated multi-omic approaches like ProteoMix can provide additional insights into pathways and processes involved in oncogenesis and how they manifest as clinical phenotypes. As CPTAC moves into its second phase and characterizes more samples across more cancer types, models such as ProteoMix may leverage this valuable data to improve understanding of the molecular basis of cancer.

## Methods

### Data Processing

Molecular data were produced from tissue bio-specimens from three cancer cohorts: breast invasive carcinoma (BRCA), colorectal adenocarcinoma (COADREAD), and ovarian serous cystadenocarcinoma (OV) (Table 1). All data used in this study comes from samples obtained from the TCGA Biospecimen Core Resource (6,9,10,24–26).

#### DNA Methylation

CpG site methylation levels/percentages were measured using Illumina Infinium Human Methylation 27k and 450k BeadChip Platforms (24–26). We limit our observations to overlapping probes or CpG sites for cancer tissues measured using both platforms, otherwise we use all available probes. The methylation level is recorded as a beta value representing a ratio of the signal/intensity from the methylated probe over the sum of both the methylated probe and the unmethylated probes. Values close to 0 indicate low levels of DNA methylation and values close to 1 indicate high levels of DNA methylation. We removed CpG sites with more than 10% missing entries across all samples and we applied 15-K Nearest Neighbor (KNN) to impute the remaining missing values, this procedure was replicated for all molecular data types. We observed significant technical sources of variation among tissue samples processed in batches using a one-way analysis, which we corrected using COMBAT (27). To reduce dimensionality of the CpG data we applied hierarchical clustering with complete linkage and a minimum average Pearson correlation of 0.4 between values. Last, we matched clusters to their corresponding genes by mapping to the closest transcriptional start sites, where one gene may relate to many CpG clusters but each CpG cluster only maps to one gene. Therefore, we limit our analysis to DNA methylation states with cis regulation effects.

#### RNA Expression

We used transcriptomic data in MethylMix produced by RNA sequencing technology (24–26). We log-transformed the RNAseq counts and replaced infinities with a non-zero low value. Similar to our DNA methylation data processing, we estimated missing values using 15-KNN and used COMBAT to correct for batch effects (27).

#### Protein Abundance

Proteomic data used in ProteoMix was provided by CPTAC (6,9,10). Participating research institutions used the following Common Data Analysis Pipeline to produce protein level measurements: First tissue samples were enzymatically digested, cutting large proteins in a sequence specific manner into smaller peptides. Peptides were fractioned using liquid chromatography to improve downstream quality before measurements using Thermo Fisher high-resolution tandem mass spectrometry (LC-MS/MS). Next, the resultant mass ladders were matched to theoretical mass ladders in the FASTA database and subsequently assigned to peptide spectra using software tools and The Reference Sequence Database. The data was then filtered to exclude peptide fragments common to more than one protein and to only include protein-identifying or unshared peptides i.e. fragments with unique sequences. Lastly peptides were mapped to a parsimonious set of genes.

The BRCA and OV workflows used iTRAQ-labeling to increase throughput, where 3 patient samples are isotopically labelled and analyzed against a common reference standard and describe relative ion intensities. Quantities are recorded after taking the log2 ratio of the abundances. Alternatively, measurement of COADREAD samples used label free mass spectrometry technology and are reported as absolute counts, which were transformed to relative quantities by taking the log2 of quantile normalized values using the limma R package (10). OV samples collected from Pacific Northwest National Laboratory and John Hopkins University were merged and corrected for batch effects using COMBAT (27).

To remove samples compromised by protein degradation we filtered samples using the QC method described by Mertins et al. (6): we calculated the standard deviation of nonnormalized protein measurements across all genes for each sample and segmented samples into groups using a two component Gaussian mixture model. Samples belong to the poor-quality group i.e. higher mean standard deviation were excluded from study. Applying this method we discarded 28 BRCA and 5 OV samples. Finally, for each cancer we removed samples with greater than 75% missing values, estimating the remaining missing values using 15-KNN algorithm (28).

### Algorithm

#### Step 1: Fit mixture model to methylation data

As described earlier methylation levels are recorded as beta values or values ranging from 0 to 1 representing the percentage of methylation and therefore gene values across all samples are beta distributed. MethylMix identifies subgroups of patients with a distinct methylation pattern or state by finding the beta mixture model with the number of components that best describe the data. To map samples to subgroups we iteratively add components requiring that each additional component improve the Bayesian Information Criterion (BIC) to avoid overfitting. To define the most descriptive subgroups we include methylation measurements across all samples, however our model integrates epigenetic data with proteomic and transcriptomic data using only the subset of these samples with available matched data (Table 1).

#### Step 2: Compare methylation to normal tissue

To identify differentially methylated CpG clusters we compare the mean methylation level - the mean value of the beta mixture component - to the mean methylation level of normal samples. To measure if an observed difference is significant we perform a Wilcoxon rank sum test with a Q-value cutoff of 0.05, using both p-value multiple testing correction with False Discover Rate (FDR). As an additional measure, we require a minimum difference of 0.10 based on the platform sensitivity (29). If significant, the difference in methylation level between the mode and normal is recorded as the Differential Methylation value or DM value for each methylation state.

#### Step 3: Select for functionally predictive genes

Next, we filter our set of genes, requiring that genes be not only differentially methylated when compared to normal but also predictive of gene expression or protein abundance. Hyper-methylation should lower gene expression and corresponding protein abundance when compared to the normally methylated samples, therefore we only accept genes that have a negative correlation between methylation level and downstream gene products. Note this assumption is only explanatory of methylation at promoter regions and does not necessarily apply to methylation at the gene-body or 3’ and 5’ untranslated regions (UTRs). To assess the likelihood that methylation events are functional, MethylMix uses the relationship between methylation and gene expression, whereas ProteoMix examines the effect on protein abundance. In both cases, we perform a linear regression between methylation levels and RNA expression or protein abundance data respectively. We use the R-square statistic to estimate the magnitude of the correlation and used cutoffs at R-square greater than 0.05 and a p-value less than 0.001.

Applying the procedures outlined above for MethylMix and ProteoMix each produces a list of candidate cancer drivers (referenced as MethylMix and ProteoMix genes) and a corresponding matrix of DM-values for identified CpG clusters across all samples. To assess the validity of each list we used orthogonal clinical and biological data to assess utility for downstream analysis and relevance to disease state.

### Evaluation

#### GO Term Enrichment

To describe the underlying biological processes captured by each model, we tested for enrichment of Gene Ontology (GO) terms in MethylMix and ProteoMix genes. This analysis was implemented using the PANTHER Classification System’s statistical overrepresentation tool (30) with the following settings: Homo-sapiens for organism, the background set to include all genes with matching protein and RNA data, and either ProteoMix or MethylMix genes for input. Enrichment was calculated using fisher’s exact test. For each gene list we rank terms using significance of the test statistic with a minimum p-value of 0.10.

#### Methylation Subtypes

With the matrices of DM values for our CpG clusters we performed consensus clustering to identify robust groupings of patients based on epigenetic signatures (31). Our analysis for each cancer cohort used the following parameters: maximum number of clusters is 6, number of bootstrap subsamples is 500 with 0.8 the proportion of the subsample, and our method uses k-means cluster algorithm and Euclidean distance. To identify the optimal number of clusters we inspected the proportion of ambiguous classification (PAC Score) (32,33), and the consensus heatmap and values, where the score/index between two samples is the proportion of clustering runs in which the two items are clustered together. We define the intra cluster consensus as the mean of all pairwise consensus scores between samples clustered in the same group, and inter cluster consensus as the mean of all consensus indexes between a sample and all the other samples clustered in different groups. A robust clustering result ideally shows high intra cluster consensus and low inter cluster consensus. We tested for association between cluster assignments and several disease features, using a Chi-squared test for categorical variables such as molecular subtype labels or a Kruskal-Wallis test for ordinal values such as tumor grade. Our analysis includes genetic, molecular, and clinical annotations, which were collected from supplementary tables from the original TCGA publications (24–26) in addition to annotations downloaded using the TCGAbiolinks R package (34).

#### Enrichment for putative tumor markers

We compared MethylMix and ProteoMix genes by investigating their enrichment in genes related to disease progression. We used correlation of gene expression with cancer stage and tumor size to identify potential genes capturing disease progression. We took the spearman correlation between gene expression levels and these clinical variables using all available samples. We selected genes using a p-value cutoff of 0.05 and biased for genes with greater likelihood of relevance by taking top 50th quantile in sample variance. Next, we filtered for only relationships that can be explained by methylation, such that genes identified as hyper-methylated in cancer tissue were required to show a negative correlation between gene expression and disease progression (tumor-suppressor genes) and hypo-methylated genes positively correlated (oncogenes). To assess each models’ likelihood in picking up genes related to disease progression we examined the overlap between these genes and the ProteoMix and MethylMix genes, using Fisher’s exact test to evaluate significance.

## Acknowledgments

Research reported in this publication was supported by National Library of Medicine Extramural Programs Grants, under award number T15-LM-007033. The funders had no role in study design, data collection and analysis, decision to publish, or preparation of the manuscript.

---

**Supplementary Table 1**. Gene level results from MethylMix and ProteoMix for each cancer site and summary statistics from linear regression taken between DNA-methylation beta values and gene-expression or protein abundance values, respectively.

**Supplementary Table 2**. Results for spearman correlations taken for each gene mRNA-protein pair.

**Supplementary Table 3**. Gene Ontology terms enrichments found for each cancer site from MethylMix and ProteoMix gene lists.

**Supplementary Table 4**. Enrichment found between cluster assignments and various clinical and molecular features, taken using Chi-squared and Kruskal-Wallis tests.

